# De-biased sparse canonical correlation for identifying cancer-related trans-regulated genes

**DOI:** 10.1101/2024.08.15.608166

**Authors:** Nathan Huey, Diptavo Dutta, Nilanjana Laha

## Abstract

In cancer multi-omic studies, identifying the effects of somatic copy number aberrations (CNA) on physically distal gene expressions (trans-associations) can potentially uncover genes critical for cancer pathogenesis. Sparse canonical correlation analysis (SCCA) has emerged as a promising method for identifying associations in high-dimensional settings, owing to its ability to aggregate weaker associations and its improved interpretability. Traditional SCCA lacks hypothesis testing capabilities, which are critical for controlling false discoveries. This limitation has recently been addressed through a bias correction technique that enables calibrated hypothesis testing. In this article, we leverage the theoretical advancements in de-biased SCCA to present a computationally efficient pipeline for multi-omics analysis. This pipeline identifies and tests associations between multi-omics data modalities in biomedical settings, such as the trans-effects of CNA on gene expression. We propose a detailed algorithm to choose the tuning parameters of de-biased SCCA. Applying this pipeline to data on estrogen receptor (ER)-associated CNAs and 10,756 gene expressions from 1,904 breast cancer patients in the METABRIC study, we identified 456 CNAs trans-associated with 256 genes. Among these, 5 genes were identified only through de-biased SCCA and not by the standard pairwise regression approach. Downstream analysis with the 256 genes revealed that these genes were overrepresented in pathways relevant to breast cancer.

## 1. Introduction

Somatic DNA variations—non-inherited changes acquired during an individual’s lifetime—play a crucial role in the development and progression of various complex diseases including cancers (Chan *et al*., 2019; Greenman *et al*., 2007). Among these variations, somatic copy number aberrations (CNAs)—small duplications or losses of DNA elements—often affect the expression of critical oncogenes and tumor suppressor genes (Taira *et al*., 2013; Steele *et al*., 2022). Recent advances in high-throughput technologies have enabled the generation of extensive genome-wide data on CNAs and gene expressions. Integrative multi-omics analyses of these data provide valuable insights into biological processes and genetic mechanisms, aiding disease management and biomarker-based therapies (Bhattacharya *et al*., 2020; Cho, 2020). In the current work, we primarily focus on identifying trans-associations between CNAs and gene expressions due to this biological interpretation in terms of actionable biomarkers. However, our pipeline can be generalized to perform integrative multi-omics analysis of any two data modalities depending on the scientific objective of the study.

Most cancer genomics research has traditionally focused on the effects of cancer-related somatic CNAs on nearby genes (cis-association/cis-effects) (Camilleri-Broet, 2012; Tamborero *et al*., 2013). In contrast, recent studies suggest that a greater proportion of disease variation is cumulatively explained by the association of DNA elements with physically distal genes (trans-associations/effects), which can highlight critical biological mechanisms and potential disease biomarkers (Liu *et al*., 2019). Current methods for exploring the association between multi-omic data in cancer genomic studies, such as CNAs and gene expressions, include pairwise association studies (Bussey *et al*., 2006; Salari *et al*., 2010; Alonso *et al*., 2017), regularized individual feature analysis (Peng *et al*., 2010; Shi *et al*., 2015; Grist *et al*., 2022), and non-linear approaches such as piecewise linear regression spline (Leday *et al*., 2013), segmented regression (Nemes *et al*., 2012), and non-linear Poisson (Jamalzadeh *et al*., 2024). However, such methods are not readily applicable to detecting trans-associations due to the lower statistical power resulting from weaker effect sizes of trans-associations and moderate sample sizes. These considerations, combined with the high dimensionality of current cancer genomic data, pose unique analytic as well as methodological challenges (Alonso *et al*., 2017; Wheeler *et al*., 2019). Therefore, there is a pressing need for specialized statistical methods to analyze such high-dimensional multi-omic data for reliable inference of trans-associations that could improve the detection of genomic biomarkers.

An emerging alternative approach is Sparse Canonical Correlation Analysis (SCCA) (Bao *et al*., 2019a; Witten *et al*., 2009; Parkhomenko *et al*., 2009; Wilms and Croux, 2015; Lee *et al*., 2011), an extension of traditional canonical correlation analysis with sparsity constraints making it amenable to be used in high dimensional setting such as cancer genomic studies. Instead of identifying potentially associated CNA-gene pairs, SCCA selects sets of CNAs and genes potentially associated with each other. The advantages of applying SCCA, especially to detect trans-associations between CNAs and genes, are two-fold: (A) SCCA aggregates multiple, potentially weaker associations across gene and CNA sets, enabling the detection of associations that may be overlooked by standard pairwise methods or sequential approaches. (B) SCCA facilitates biological interpretation by selecting smaller sets of genes and CNAs, through sparse linear combinations, where the selected gene-set represents the genes regulated by the CNAs selected in the corresponding CNA-set. This is particularly amenable to interpretation since it is well known that genes act as a network rather than in isolation, and none of the previously mentioned methods attempt to uncover such gene-set structure. In fact, SCCA has successfully been used to identify the trans-regulation of gene sets by germline variants in various complex diseases and to investigate the association between somatic variation and gene expression (both cis and trans) in breast cancer (Dutta *et al*., 2022a,b).

However, the current implementation and standard approaches using SCCA suffer from a notable limitation, which is non-negligible in the context of trans-association. The weaker effect sizes of trans-associations and the high dimensionality of CNA and gene expression data can often lead to false detections (Alonso *et al*., 2017). Standard false discovery control methods such as FDR (false discovery rate) control and Bonferroni correction can only be applied if hypotheses on a gene expression’s associations with distal CNAs are testable (Benjamini and Hochberg, 1995; Benjamini *et al*., 2001). In the context of CCA, this translates to testing whether a quantity called the CCA “loading” is zero for this gene expression. These tests substantially differ from testing whether the canonical correlations themselves are zero. The latter tests, although well-studied, only inform whether the selected gene expressions as a set are associated with the selected CNA set (Witten *et al*., 2009; McKeague and Zhang, 2022; Senar *et al*., 2024; Bao *et al*., 2019b). Most mainstream SCCA methods, such as those by Mai and Zhang (2019), Gao *et al*. (2017), and Witten *et al*. (2009), use *ℓ*_1_ regularization, leading to asymptotic bias in the SCCA loading estimators (Leeb and Pötscher, 2005, 2006, 2008; Pötscher and Leeb, 2009). These estimators have intractable asymptotic distributions and are not suitable for 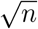-level asymptotic tests, complicating false discovery control (Kessler and Levina, 2023; Laha *et al*., 2023). Although these tests can be conducted using re-sampling, suitable re-sampling methods for the high-dimensional setting considered here are currently unavailable (see Section 4 for more details).

In our recent work Laha *et al*. (2023), we developed a de-biasing procedure for SCCA estimators. De-biasing is a popular technique for correcting the *ℓ*_1_-regularization-induced asymptotic bias in high dimensional estimation (Javanmard and Montanari, 2014; van de Geer *et al*., 2014; Zhang and Zhang, 2014; Janková and van de Geer, 2021). De-biased SCCA offers two main advantages: (1) Its loadings are asymptotically Gaussian, enabling valid z-tests and standard false discovery control methods. (2) De-biased SCCA can be combined with SCCA methods such as Mai and Zhang (2019)’s SCCA, which account for correlations within variable sets, unlike Witten *et al*. (2009); Parkhomenko *et al*. (2009)’s SCCA. This is important for our particular application, as we will explain.

In this study, we integrate the de-biasing method with Mai and Zhang (2019)’s SCCA to identify trans-associated genes with breast cancer-related CNAs using data from 1904 breast cancer patients from the METABRIC study (Pereira *et al*., 2016). In particular, we focus on estrogen receptor (ER) status which indicates whether the tumor cells have receptors that can bind to estrogen and is one of the key predictors of cancer aggressiveness and prognosis. This study is one of the first attempts to leverage the recent theoretical advancements in inferential SCCA to produce biologically plausible results from cancer genomic data through a comprehensive pipeline using de-biased SCCA. Although Laha *et al*. (2023) demonstrated the theoretical efficacy of the de-biasing method and provided numerical evidence of its success in controlled settings, its application to real-world biomedical data presents several challenges. Our proposed approach consists of three key steps: (1) Identifying CNAs associated with estrogen receptor (ER) status using Minimum Redundancy Maximum Relevance (MRMR), a supervised variable selection technique. (2) Identifying topologically independent gene modules representing major gene expression clusters among the distal genes (distal to the ER-related CNAs identified in step 1). (3) Using de-biased SCCA to identify genes trans-regulated by ER-related CNAs. In the following sections, we first introduce the description of the METABRIC dataset and then the statistical methodology. De-biased SCCA depends on multiple tuning parameters and is sensitive to their selection. Therefore we introduce tailored algorithms to address this. We then discuss the results of applying de-biased SCCA on METABRIC data and demonstrate its advantages over the regression-based pairwise association method.

## 2. Methods

### 2.1 The dataset

Our data is from the Molecular Taxonomy of Breast Cancer International Consortium (METABRIC), one of the largest publicly available sources of comprehensive molecular characterization of breast cancer patients (Pereira *et al*., 2016). The current version includes detailed sample characteristics, recorded clinical outcomes, and molecular profiling from 2509 patients. Among them, 1904 patients had complete information on ER status, 24,368 gene expressions, and 22,544 CNAs. CNA sites take values of {−2, −1, 0, 1, 2}, where negative values indicate the deletion of base pairs on one or both chromosomes, respectively, while positive values of CNA indicate an insertion. As mentioned above, we first identified a set of 500 CNAs associated with individual-level ER status through MRMR. Given we aim to identify genes trans-associated with this set of 500 CNAs, we then excluded genes with a transcription start site within a 1Mb neighborhood of the selected CNAs, resulting in 10,756 genes. This set of 500 ER-related CNAs and 10,756 genes distal to each of the CNAs were considered for our analysis.

### 2.2 Statistical method

This section presents an overview of SCCA and de-biased SCCA. Given our scientific goal of identifying trans-associations between CNAs and gene expression, we explain the methods in terms of these two data modalities, but can easily be generalized to any pair of omics data. Suppose we have individual-level data from *n* patients. Further suppose, for each patient, we have CNA data at *p* sites and the expression level of *q* genes. For *i* = 1, …, *n* and *j* = 1, …, *p*, let *X*_*ij*_ denote the number of gene insertions or deletions at the *j*th CNA site for patient *i*. We let **X** denote the corresponding CNA matrix. Similarly, we let **Y** be the matrix of normalized gene expression levels for *q* genes across *n* individuals, i.e., *Y*_*ij*_ is the normalized expression level of the *j*th gene in the *i*th individual where *i* = 1, …, *n* and *j* = 1, …, *q*. Also, letting 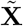 and 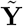 denote the centered and scaled versions of **X** and **Y**, respectively, we define the sample covariance matrices as 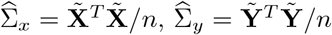, and 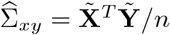. The population versions of these matrices will be denoted by Σ_*x*_, Σ_*y*_, and Σ_*xy*_, respectively. The operator norm of any matrix *A* will be denoted by |*A*|_*op*_. The *l*_1_ norm of any vector *x* will be denoted by |*x*|_1_. For any integer *k, I*_*k*_ will denote the identity matrix of order *k*.

### 2.3 Sparse Canonical Correlation Analysis (SCCA)

Current genetic studies hypothesize that a large majority of genetic effects on complex diseases, such as breast cancer, are mediated by cascading trans-effects on a few “core genes” (Liu *et al*., 2019). In such cases, SCCA can be conceptualized as a method to select small groups of CNAs and gene expressions that are highly correlated. SCCA aims to find the sparse linear combination of CNAs (*α* ∈ ℝ^*p*^: CNA component) and gene expressions (*β* ∈ ℝ^*q*^: gene-expression component) that maximize the correlation between **X***α* and **Y***β*. Most mainstream SCCA implementations use *l*_1_ regularization to enforce sparsity on *α* and *β* (Mai and Zhang, 2019; Witten *et al*., 2009; Gao *et al*., 2017). The corresponding maximization problem is given by

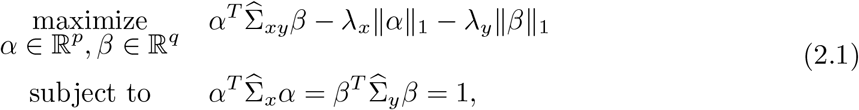

where ∥ · ∥_1_ is the *l*_1_ norm and *λ*_*x*_, and *λ*_*y*_ are two positive tuning parameters controlling the sparsity level. The solutions to (2.1) are referred to as the leading canonical directions, which we denote by 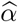 and 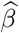, respectively, and are termed loading vectors. The population version of 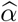 and 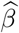 will be denoted by *α*_0_ and *β*_0_, respectively. The *l*_1_-penalty on *α* and *β* in (2.1) renders *α* and *β* to be sparse. Values of tuning parameters *λ*_*x*_ (or *λ*_*y*_) control the sparsity of 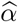 (or 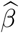). The biological interpretation of 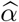 (and 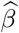) is that the non-zero elements represent the set of CNAs (and genes) selected by SCCA. Thus if 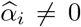, it generates the hypothesis that the *i*^*th*^ CNA is associated with the corresponding genes selected in 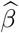 and vice versa. The maximum value attained by the optimization program (2.1) is known as the leading canonical correlation, and will be denoted by 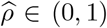. It represents the strength of the overall association between the selected sets of CNAs and genes. The population version of 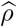 is 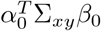, which will be denoted by *ρ*_0_ from now on. *ρ*_0_ *>* 0 unless the genes and the CNAs are completely uncorrelated, in which case *ρ*_0_ = 0. In this scenario, *α*_0_ and *β*_0_ will be zero vectors.

The optimization problem in (2.1) is non-convex and hard to optimize due to the non-convex elliptical constraints 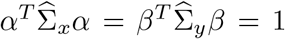. Existing approaches use clever convex relaxations of the constraints (Gao *et al*., 2017) or iterative programs to solve (2.1) (Mai and Zhang, 2019; Wilms and Croux, 2015). We use Mai and Zhang (2019)’s method to obtain preliminary SCCA estimators 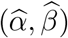, which subsequently undergo bias correction by Laha *et al*. (2023)’s de-biasing method. From now on, we will refer to Mai and Zhang (2019)’s estimator as the SCCA-MZ estimator. We preferred SCCA-MZ over COLAR based on a pilot comparative analysis, which revealed SCCA-MZ to be more robust to false positives than COLAR (see Supplement B.2). Additionally, Laha and Mukherjee (2022)’s simulations show that this estimator has impressive variable selection properties under controlled settings. Tuning parameter selection, including that of *λ*_*x*_ and *λ*_*y*_, will be discussed in Section 2.5.

We will clarify an important point here. A popular SCCA approach, pioneered by Witten *et al*. (2009); Solari *et al*. (2019); Parkhomenko *et al*. (2009) among others, replaces the constraint 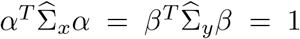 with *α*^*T*^ *D*_1_*α* = 1 and *β*^*T*^ *D*_2_*β* = 1, where *D*_1_ and *D*_2_ are diagonal matrices, thus assuming independence between CNAs and genes in the current context. While this approach gains computational efficiency, it ignores correlations within genes and CNAs, which might not be realistic or conducive to biological interpretation. In particular, if a CNA and a gene are associated, SCCA loadings, by assuming independence between genes, will tend to select a broader set of genes all correlated with the true CNA-associated gene, a tendency also observed by Mai and Zhang (2019). This diverges from our main scientific goal, in that by accounting for correlations, we want to identify independent sources of trans-association that are potentially causal, and not merely arising due to correlation with the actual causal variable.

### 2.4 De-biasing the SCCA estimator

SCCA estimators such as SCCA-MZ incur a first-order bias due to the use of *l*_1_ penalty and lack a tractable asymptotic distribution (Kessler and Levina, 2023). In this section, we will discuss how the de-biasing method can be used to conduct hypothesis tests with such SCCA estimators.

Our hypotheses of interest, i.e., that the *i*th CNA has no association with the genes is equivalent to *H*_0_: (*α*_0_)_*i*_ = 0, while the hypothesis that the *j*th gene has no association with the CNAs is equivalent to *H*_0_: (*β*_0_)_*j*_ = 0 (Gao *et al*., 2017; Anderson, 1962). Note that testing *H*_0_: (*α*_0_)_*i*_ = 0 or *H*_0_: (*β*_0_)_*j*_ = 0 is equivalent to testing 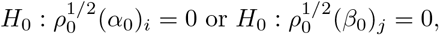, respectively. The latter versions of the hypotheses have some advantages, as described below. Laha *et al*. (2023) showed that 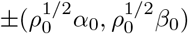 are the unique minimizers of the unconstrained minimization problem:

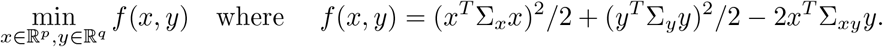

Using the above representation, they show that 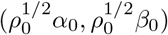 can be efficiently estimated via a Newton-Raphson one-step bias correction starting with the preliminary estimator 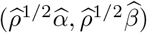. The de-biasing procedure uses the scaled version 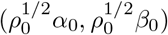 instead of (*α*_0_, *β*_0_) because no similarly useful representation is known for the latter.

The de-biased estimators 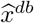 and 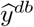 of 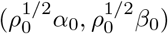 are given by

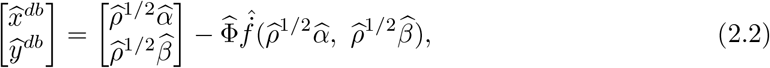

where 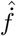 is an estimator of the gradient of *f*, and 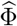 is an estimator of the inverse of the Hessian matrix of *f* at 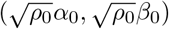. The vector 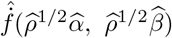 can be estimated straightforwardly using a plug-in estimator; see Laha *et al*. (2023) for details. Constructing 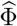, however, involves inverting the Hessian of *f*, a high-dimensional matrix. Laha *et al*. (2023) recommends using the nodewise lasso algorithm of Van de Geer *et al*. (2014). This algorithm estimates each column of the Hessian inverse separately through lasso regressions. The *ℓ*_1_ penalty of these lasso regressions induces sparsity in the columns of 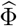. The sparsity level is controlled by a common penalty parameter *λ*_*NL*_.

Laha *et al*. (2023) showed that, under mild regularity conditions, the de-biased estimators defined in (2.2) are elementwise asymptotically normal. Specifically, for each *i* ∈ {1, …, *p*} and *j* ∈ {1, …, *q*}, it holds that 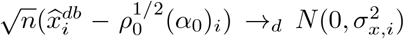 and 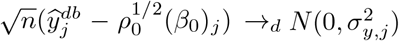, where 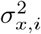 and 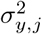are the asymptotic variances of 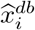 and 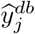,respectively. Under the null hypothesis that the *j*th gene has no association with the CNAs, i.e., (*β*_0_)_*j*_ = 0, the Z-score is defined as

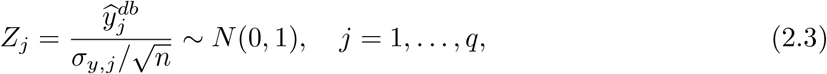

which can be used to construct a Z-test. In this test, we replace *σ*_*y,j*_ by its consistent estimator provided by Laha *et al*. (2023). Similarly, we can construct Z-scores for the CNAs, leading to analogous Z-tests. Consequently, *p* + *q* hypothesis tests can be conducted for each gene and each CNA. The combined type I error of these *p* + *q* many tests can be high, requiring multiple hypothesis testing correction for controlling the false discovery rate (Benjamini and Hochberg, 1995; Benjamini *et al*., 2001). Laha *et al*. (2023) showed that the marginal distribution functions of the *p* + *q* Z-scores converge uniformly when *p, q*, and *n* are large. This result enables the application of standard false discovery rate control methods on the p-values of the *p* + *q* Z-tests resulting from (2.3). In particular, we use the method of Benjamini Hochberg (Benjamini and Hochberg, 1995).

### 2.5 Tuning

As indicated in Sections 2.3 and 2.4, our method involves three main tuning parameters, *λ*_*x*_, *λ*_*y*_, and *λ*_*NL*_. These parameters are penalty parameters associated with *l*_1_-regularization, controlling the sparsity levels of 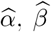 and 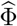, respectively. Below, we describe our procedure for selecting these tuning parameters, outlined in Algorithm 1. We first discuss the tuning parameters *λ*_*x*_ and *λ*_*y*_ for SCCA-MZ, as the tuning of *λ*_*NL*_ depends on them.

#### SCCA-MZ tuning

Following Mai and Zhang (2019)’s recommendations, we choose the tuning parameter combination that maximizes the resulting estimated canonical correlation. By considering a two-dimensional grid of (*λ*_*x*_, *λ*_*y*_) values, we estimate the canonical correlation for each tuning parameter combination using 5-fold cross-validation. See Algorithm E.1 in Supplement E for more details. We denote the selected tuning parameters at the end of this stage by 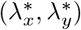, and the corresponding SCCA-MZ estimators by 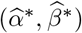.

#### Nodewise lasso tuning

To tune the nodewise lasso parameter *λ*_*NL*_, we split the dataset into two roughly equal parts *S*_1_ and *S*_2_. First, using *S*_1_ as the training set, we compute the inverse Hessian estimator 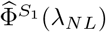 for each value of the tuning parameter *λ*_*NL*_ on a grid. This estimator is computed following the nodewise lasso algorithm (Algorithm 2 in the Supplement of Laha *et al*., 2023). This algorithm uses the SCCA-MZ estimators 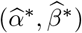 discussed above. Next, using *S*_2_ as the test set, we compute the estimator 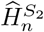 of the Hessian of *f* at 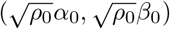 using Algorithm 2. For each value of the candidate tuning parameter *λ*_*NL*_, we calculate a loss as follows:

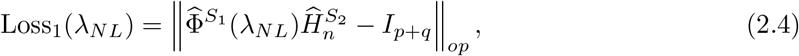

where *I*_*p*+*q*_ is the identity matrix of order *p* + *q*. Next, we swap the roles of *S*_1_ and *S*_2_, leading to a second loss:

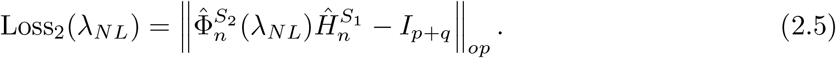

The final loss for *λ*_*NL*_ is then defined as the sum of the losses in (2.4) and (2.5):

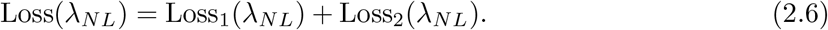

We compute Loss(*λ*_*NL*_) for each candidate nodewise lasso tuning parameter, and the final node-wise lasso tuning parameter, 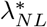, is set to the value yielding the minimum loss. Observe that we estimate the Hessian using a dataset different from the one used to estimate 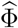. Simulations suggest that using the entire dataset to estimate the Hessian tends to select higher values of *λ*_*NL*_, and the resulting losses are also higher.

The computation of the Hessian requires estimators of Σ_*x*_, Σ_*y*_, and Σ_*xy*_. To this end, we use the Ledoit-Wolf Linear Shrinkage estimator of covariance matrices (Ledoit and Wolf, 2004) instead of the sample covariance matrices. Pilot simulations indicated that the Ledoit-Wolf Linear Shrinkage estimators provide better stability for this Hessian estimator. Notably, the nodewise lasso estimator also relies on estimators of Σ_*x*_, Σ_*y*_, and Σ_*xy*_. However, similar to Laha *et al*. (2023), we use the sample covariance matrices in the nodewise lasso estimator as the asymptotic normality of the scaled loadings specifically requires the sample covariance matrices at this step. The pseudocode for the nodewise lasso tuning procedure is provided in Procedure “Nodewise Lasso Tuning” in Algorithm 1.

## 3. Results: Analysis of METABRIC data

### 3.1 Supervised selection of ER-related CNAs

We applied the de-biased SCCA as described in Section 2.4 on METABRIC data (see Section 2.1) to identify genes that are trans-regulated by CNAs associated with estrogen receptor (ER) status. We first selected a set of 500 CNA sites associated with ER status agnostic of the gene expressions. Thus, this screening does not alter the subsequent analysis to identify associations between CNAs and genes. Hence, this step does not affect the type I error of any downstream inference using SCCA. The rationale for this step is to identify key CNA sites that influence ER status. Any gene which is identified to be trans-associated with these CNAs can be interpreted as a downstream target gene via which the effects of these CNAs might be cascaded.

For selecting the ER-associated CNAs, we used MRMR, a supervised feature selection method widely used in genomic data analysis (Bugata and Drotar, 2020; Ding and Peng, 2005; Chandrashekar and Sahin, 2014; Tadist *et al*., 2019), including analyses with the METABRIC dataset (Mucaki *et al*., 2016). MRMR is a forward selection method, iteratively adding features to a predictive set with maximum association to the response, while ensuring minimal redundancy among the selected features. As in Bugata and Drotar (2020), we used the mutual information to measure the association between variables. Of particular relevance to us, MRMR also returns a ranking of the features. Although there are many other supervised feature selection methods, such as *l*_1_-penalized logistic regression (Genç, 2022), Feature Ordering by Conditional Independence (FOCI) (Azadkia and Chatterjee, 2021), and RELIEF (Robnik-Šikonja and Kononenko, 2003), a preliminary comparison of these methods revealed MRMR to be more stable on our dataset compared to these other methods. See Supplement A for more details on this analysis.

We used an ensembling approach to increase the stability of the MRMR feature selection procedure. To this end, we first generated 40 bootstrap samples following Abeel *et al*. (2010), who used a similar approach for identifying cancer biomarkers in a microarray dataset. We then used the MRMR algorithm on each bootstrap sample to select the top 500 CNAs. This led to 40 sets of top 500 CNAs – we call these the “bootstrap selection sets”. The final set of 500 CNA sites was then selected based on the frequency of occurrence in these 40 bootstrap selection sets. To illustrate: any CNA site found in all 40 bootstrap selection sets was first added to the final set. Then any CNA sites found in exactly 39 of the bootstrap selection sets were added to the final set, and so on. This continued until we reached the level *n* such that the total number of CNAs in the final selection set would exceed 500 if all the CNAs in exactly *n* selection sets were added to the final set. We describe this situation as a “tie”. We used the CNA rankings returned by the MRMR algorithm to break the ties. Specifically, the tie-break was decided by the average ranking (across the 40 bootsrap samples) of the CNAs found in exactly *n* samples.

### 3.2 Gene Set clustering

Given the 500 ER-related CNAs identified in the previous step, we then extracted the expressions of all genes located at least 1 Mb away from these CNAs, resulting in 10,756 genes. However, joint analyses of unrelated and potentially independent genes can result in the loss of biological interpretation of the identified gene-sets. Hence, we sought to identify major clusters of genes through co-expression analysis, agnostic of the CNAs. We used weighted gene co-expression network analysis (WGCNA) to build gene co-expression networks (Langfelder and Horvath, 2008; Horvath, 2011). WGCNA is a widely used method in genomic data analysis (Horvath, 2011). It uses hierarchical clustering to partition a gene set into distinct modules with highly correlated expression patterns. The idea is that because genes with similar expression patterns are clustered together, WGCNA modules will represent sets of genes with related functions.

The optimal number of clusters was determined via the Unweighted Pair Group Method with Arithmetic Mean (UPGMA), a default choice for the R package WGCNA (Sokal and Michener, 1958). WGCNA also automatically identifies and eliminates outliers. Eleven individuals were identified as expression outliers and excluded from further analyses. The WGCNA analysis resulted in 6 co-expression modules, with sizes ranging from 283 to 1536 genes, and 1 residual module (module 0) with 5,722 genes. Table 1 displays the number of genes in each module. The estimated canonical correlations based on the SCCA-MZ estimator were 0.90, 0.83, 0.90, 0.87, 0.92, and 0.87 for Modules 0, 1, 2, 3, 4, 5, and 6, respectively.

**Table 1.** Details of the WGCNA gene-modules. The column “Size” indicates the number of genes in each module. “Discovered” shows the number of significant genes after FDR correction at 5% level, and “Novel” lists the number of novel genes in each module. Most discovered and novel genes are from module 1.

| Module | Size | Discovered | Novel |
| --- | --- | --- | --- |
| 0 | 5722 | 10 | 0 |
| 1 | 1536 | 109 | 4 |
| 2 | 1506 | 16 | 0 |
| 3 | 830 | 9 | 0 |
| 4 | 582 | 19 | 0 |
| 5 | 297 | 27 | 1 |
| 6 | 283 | 23 | 0 |

### 3.3 De-biased SCCA analysis: METABRIC data

For each gene module, we used SCCA-MZ to obtain initial sparse estimates of the loadings for CNA and gene components. Next, we applied the de-biasing procedure described in Section 2.4 to these SCCA-MZ estimators for each gene module separately. The de-biased estimators were subsequently used for testing the trans-association between the selected CNAs and the gene sets. We corrected for false discovery rate due to multiple testing by applying the Benjamini-Hochberg (BH) method on the p-values derived from all modules, as discussed in Section 2.4.

For all modules, we tuned the nodewise lasso parameter *λ*_*NL*_ according to Algorithm 1. We used a grid of five points evenly spaced from 0.01 to 10. This grid was informed by simulations detailed in Supplement D. For tuning the parameters for the SCCA-MZ estimators in modules 1-6, we applied the SCCA-MZ tuning procedure described in Section 2.5 (Algorithm E.1 in Supplement E) with a 10×10 grid ranging from 0 to 0.10. This grid selection was guided by pilot simulation studies detailed in Supplement C. Module 0, being significantly larger, required a different approach for SCCA-MZ tuning due to its size (Table 2). As the signal-to-noise ratio diminishes with dimension, de-biased-SCCA can be potentially conservative in large modules. Using the above-mentioned grid, SCCA-MZ TUNING Algorithm selected the conservative SCCAMZ tuning parameters (0.1, 0.1) for module 0, resulting in an exceedingly sparse gene-loading vector 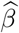 (26 non-zero elements) not amenable to any further meaningful analysis. Conversely, initiating with a very small penalty parameter (*λ*_*x*_, *λ*_*y*_) = (0.001, 0.001) renders 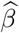 dense (955 non-zero elements), posing a risk of false positives. We chose (*λ*_*x*_, *λ*_*y*_) to be (0.01, 0.01), which yields a moderately dense gene loading vector 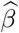 with 578 non-zero elements which is a reasonable number to perform subsequent de-biasing and hypothesis testing.

**Table 2.**
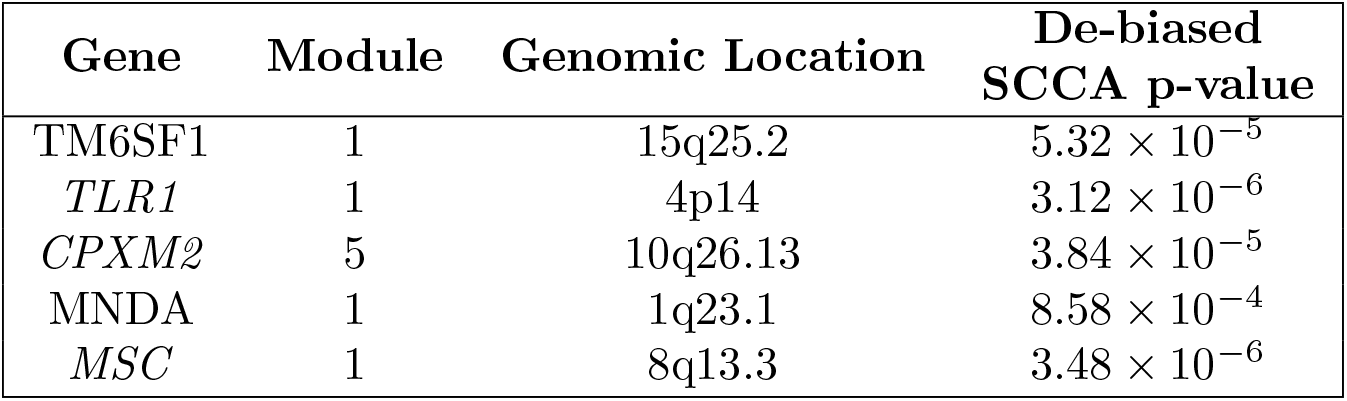
Novel Genes identified by de-biased SCCA. This table presents details on the five genes identified by de-biased SCCA, but not by the pairwise linear regression analysis. The “Module” column indicates the WGCNA module to which each gene belongs (Section 3.2). The provided genomic location of the novel genes follows the chromosomal banding system, representing the chromosome number, arm, and specific band or subregion of the gene. For instance, “15q25.2” denotes the long arm (q) of chromosome 15, subregion 25.2. The “De-biased SCCA p-value” column displays the unadjusted p-value of the Z-tests in (2.3).

| Gene | Module | Genomic Location | De-biased<br>SCCA p-value |
| --- | --- | --- | --- |
| TM6SF1 | 1 | 15q25.2 | $5.32 \times 10^{-5}$ |
| *TLR1* | 1 | 4p14 | $3.12 \times 10^{-6}$ |
| *CPXM2* | 5 | 10q26.13 | $3.84 \times 10^{-5}$ |
| MNDA | 1 | 1q23.1 | $8.58 \times 10^{-4}$ |
| *MSC* | 1 | 8q13.3 | $3.48 \times 10^{-6}$ |

### 3.4 Significant trans-associated genes

At a false discovery rate of 5%, our de-biased SCCA procedure identified 456 significant CNA sites trans-associated with 213 significant genes. Following the previous discussion, this set of 213 genes can be interpreted as the set of genes trans-regulated by the corresponding set of CNAs and potentially cascade the effects of the ER-related CNAs. Figure 1 visually represents a subset of the selected CNAs and genes. Several of the identified genes have been previously linked to breast cancer, particularly in relation to estrogen receptor activity. For example, among the 213 significant genes, we identified *ESR1*, which is crucial for mediating the effects of estrogen in various tissues, including those of the reproductive system (Chen *et al*., 2022). *ESR1* plays a significant role in breast tissue and breast cancer development, influencing cell proliferation and tumor growth, making it a primary target for hormone therapies designed to block estrogen signaling (Lei *et al*., 2019; Brett *et al*., 2021). Another example is the *FOXA1* gene, which plays a crucial role in breast cancer by modulating estrogen receptor (ER) activity and chromatin accessibility, thereby influencing gene transcription (Seachrist *et al*., 2021; Fu *et al*., 2019). Dysregulation of *FOXA1* through amplification, mutation, or altered cofactor interactions can lead to changes in ER transcriptional programs. Moreover, within these 213 genes, we detected 14 tier-1 cancer genes as per the COSMIC cancer gene census (Sondka *et al*., 2018) along with at least 35 candidate cancer driver genes overall, including *CASZ1* and *FYN*. These genes have been shown to be crucial for cell proliferation in breast cancer tumors through CRISPR knock-off experiments (Kim *et al*., 2023; Tuano *et al*., 2023).

**Fig. 1.**
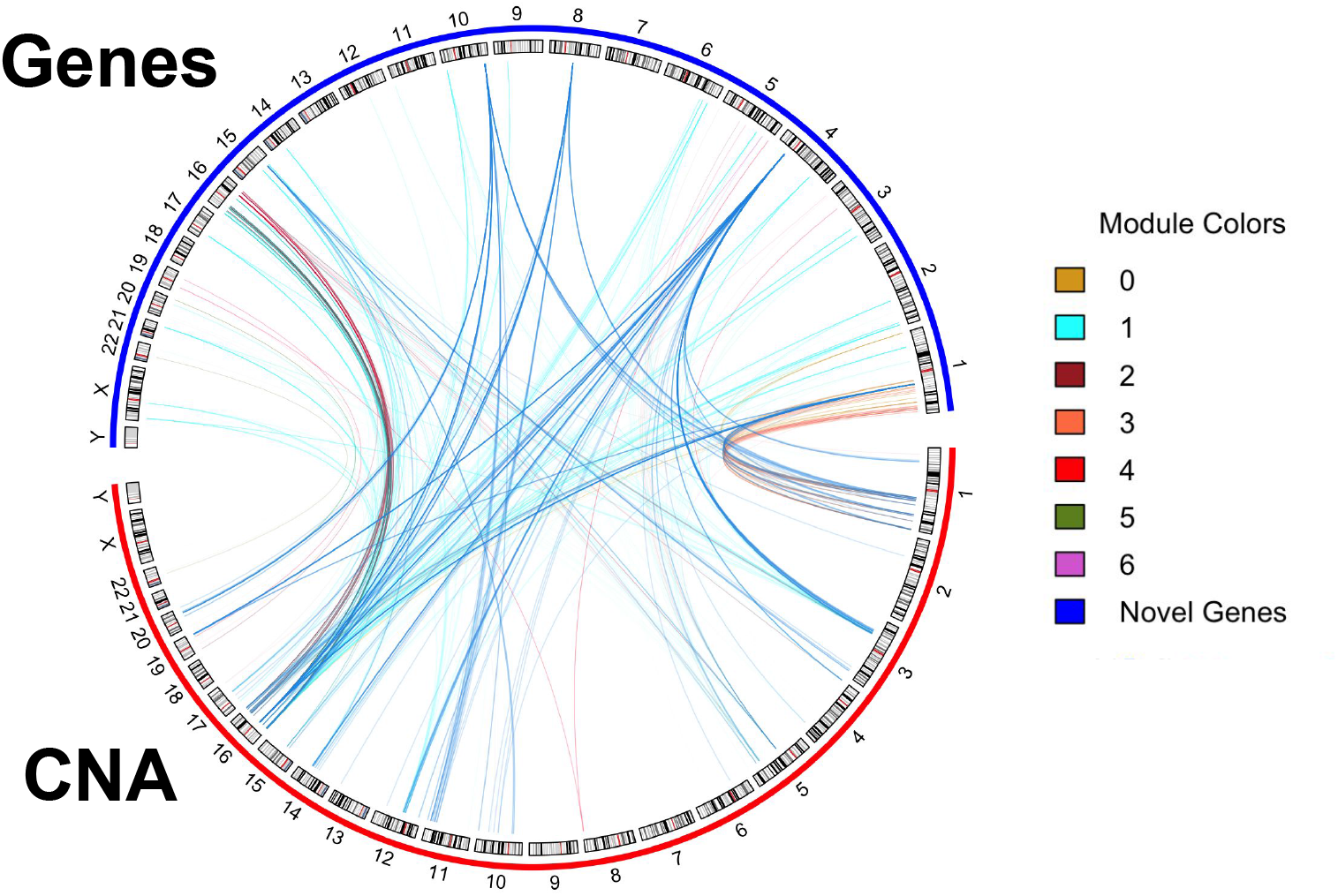
Subset of selected CNA and genes discovered via de-biased SCCA. The Circos plot illustrates the associations between the genes and CNAs detected by the de-biased-SCCA method. CNAs are represented on the bottom half, while genes are on the top half. The dark blue lines depict the five novel genes. A blue line connects a novel gene to a CNA if the corresponding pairwise linear regression p-value is below 5 *×* 10^−4^. For other gene-CNA pairs, a line is drawn if their linear regression p-value is less than 10^−30^.

As mentioned earlier in Section 1, one major advantage of the canonical correlation-based approach over standard pairwise association studies is its ability to aggregate multiple weaker associations. To illustrate this, we conducted a pairwise linear regression analysis on the identified 456 CNAs and 213 genes, followed by the standard Bonferroni correction. The heatmap in Figure 2 shows the regression p-values for all genes and CNAs. Interestingly, our de-biased SCCA identified 5 genes that were not identified using pairwise linear regression as pointed out in Figure 2. For the purpose of this manuscript, we refer to these genes as “novel” genes (See Table 2). The novel genes lacked strong trans-association with any single CNA, as shown in the heatmap in Figure 2, and were only identified by aggregating weaker associations. In contrast, the remaining 208 genes displayed strong trans-association with at least one CNA cluster. Thus, de-biased-SCCA’s detection of the five novel genes may be attributable to its ability to aggregate weaker signals.

**Fig. 2.**
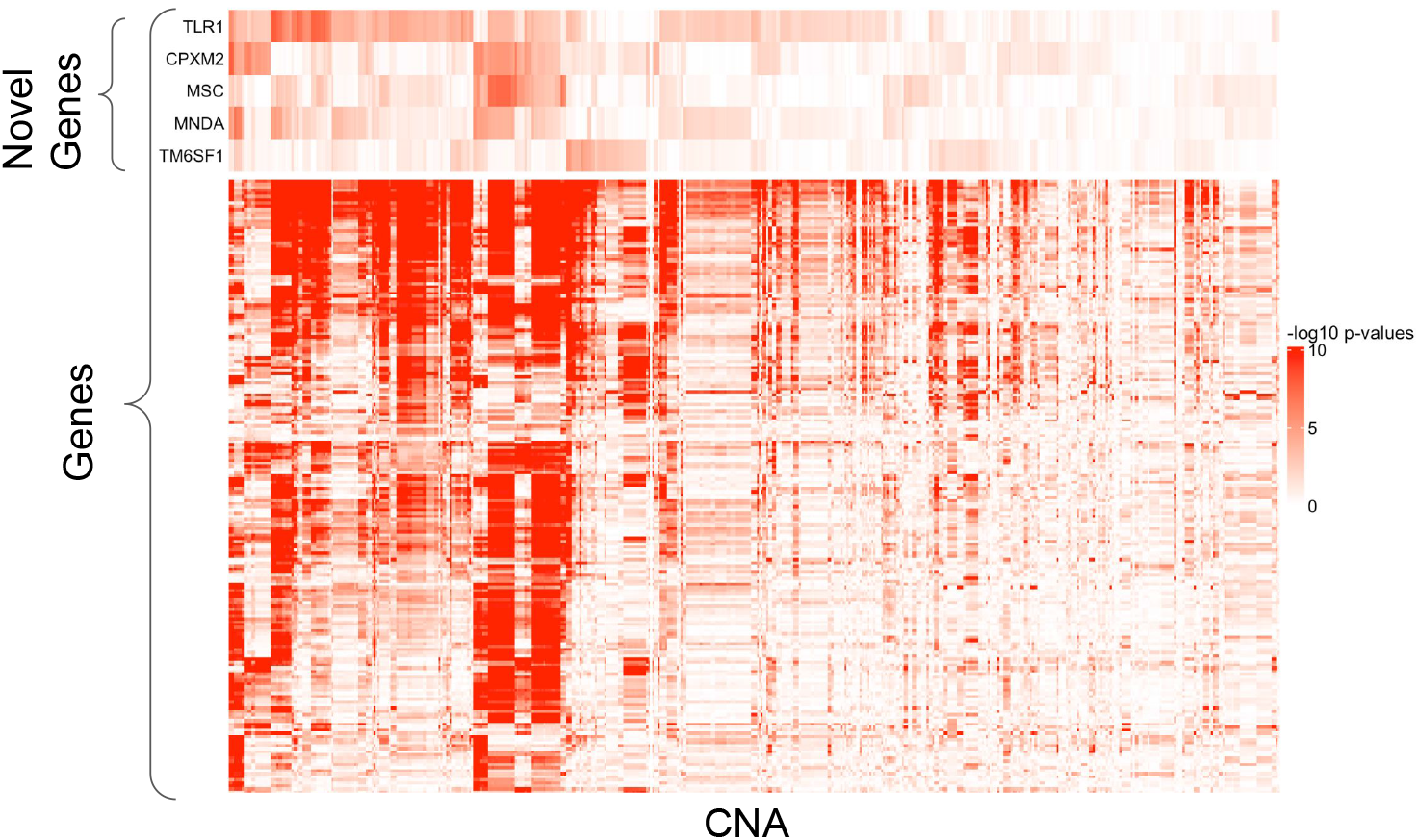
Heatmap of − log_10_ p-values from pairwise linear regression of gene expressions on CNAs. The Y-axis represents the 213 genes selected via de-biased SCCA, with the 5 novel genes highlighted at the top. The X-axis corresponds to the 456 CNAs detected by the de-biased SCCA. The p-values in this plot are the unadjusted p-values from the linear regression of each gene expression on each CNA. A color gradient from white to red indicates decreasing p-values, with white indicating lower significance and red indicating higher significance. Unlike the other 208 genes, the novel genes lack deep red regions, suggesting their lack of strong association with any single CNA cluster.

Several of the “novel” genes have been previously linked to cancer, especially breast cancer. For instance, *TM6SF1* (with a p-value of 5.32 *×* 10^−5^, Table 2) has been associated with hypermethylation in various cancers, including breast cancer (de Groot *et al*., 2016). Hypermethylation is a hallmark of cancer development as it suppresses gene expression, making *TM6SF1* a potential biomarker for early breast cancer detection (Tan *et al*., 2018). We further validated *TM6SF1* ‘s significance by examining its expression levels in cancer tissues from the Breast Invasive Carcinoma (BRCA) dataset sourced from the Cancer Genome Atlas (TCGA) repository (Lingle *et al*., 2016). In this dataset, *TM6SF1* showed significantly lower expression in tumor tissues compared to normal tissues and was nominally associated with disease-specific survival (*p*-value = 0.02; see Supplementary Figure 3). The set of our five novel genes also includes *MNDA* (p-value: 8.58*×*10^−4^, Table 2). This gene has been proposed as a master regulator in breast carcinomas and has a critical immunoregulatory function in different cancers (Tang *et al*., 2022; Briggs *et al*., 1994; Baca-López *et al*., 2012). It is anticipated that this gene potentially impacts immune invasion and tumor microenvironment (Baca-López *et al*., 2012). In the TCGA-BRCA dataset, tumor samples had a significantly higher expression of *MNDA* (see Supplementary Figure 3). Additionally, we found that point mutations in the 213 genes identified by de-biased SCCA were overall significantly associated with the abundance of CD4 and B cells in the TCGA-BRCA dataset. Both CD4 and B cells have been identified to play crucial roles in breast cancer tumor microenvironment (Zhang *et al*., 2023; Tay *et al*., 2021).

### 3.5 Gene Network Enrichment Analysis

Pathway enrichment analysis aims to identify whether a particular set of genes is over-represented or enriched among a pre-curated list of genes that constitute a biological process, mechanism, or pathway. By identifying significantly enriched pathways, this analysis can highlight the biological mechanisms that the genes might be involved in. We used the pathway enrichment analysis tool ShinyGO 0.80 (Ge *et al*., 2020) to investigate the pathways enriched for the 213 genes detected by de-biased SCCA. Figure 3 presents the enriched pathways and their significance levels. Many of these identified pathways are relevant in cancer. In particular, the identified pathways include “Schuetz breast cancer ductal invasive up” (Schuetz *et al*., 2006, FDR = 5.7 *×* 10^−26^), “Genes down-regulated in luminal-like breast cancer cell lines compared to the mesenchymal-like ones” (Charafe-Jauffret *et al*., 2006, FDR = 9.2 *×* 10^−5^), “Yang breast cancer *ESR1* bulk up” (Yang *et al*., 2006, FDR = 1.8 *×* 10^−6^), “Epithelial Mesenchymal transition” (Georgakopoulos-Soares *et al*., 2020, FDR = 2.6 *×* 10^−5^), and “KRAS signaling” (Kim *et al*., 2021, FDR = 9.8 *×* 10^−5^), all of which have established relevance in cancer. Additionally, we identified enrichment of the identified genes among targets of known cancer and immune-related transcription factors such as *SP1* and *VSX2* (Beishline and Azizkhan-Clifford, 2015; Zou and Levine, 2012). Interestingly, several novel genes were found in the above-mentioned pathways, such as *TM6SF1* and *MNDA* in the “Schuetz breast cancer ductal invasive up” pathway. These results provide further validation and independent lines of evidence for the biological relevance of the genes detected through de-biased SCCA, including the novel genes.

**Fig. 3.**
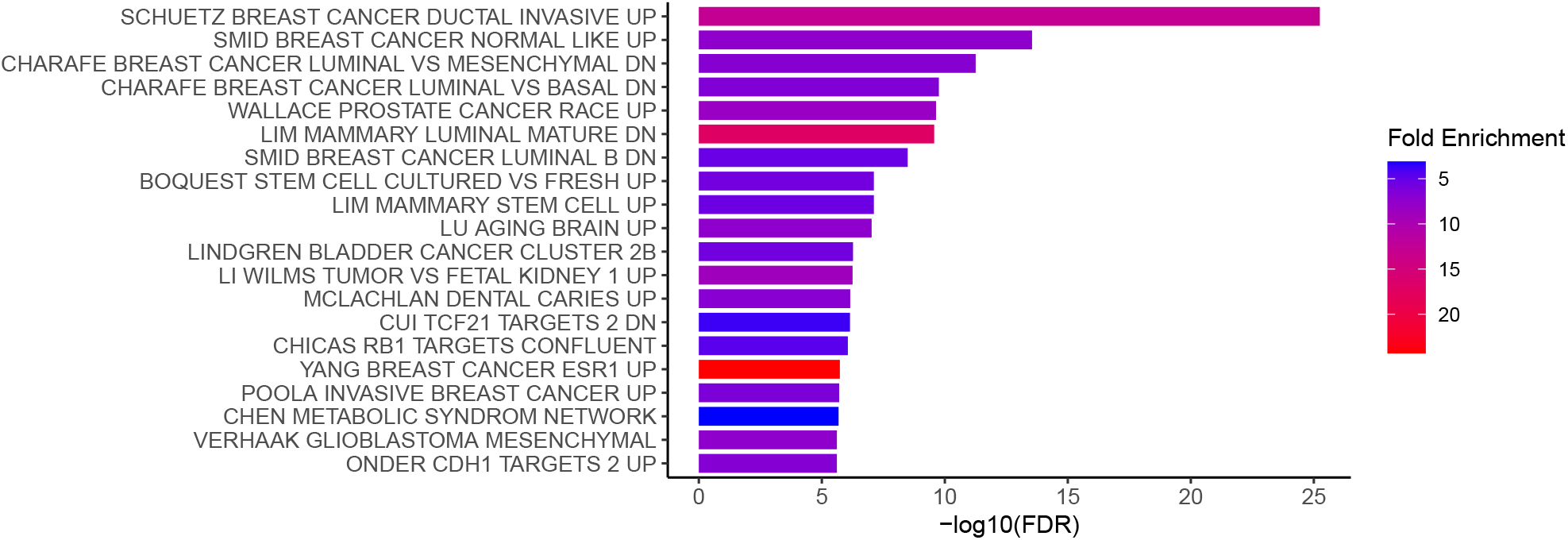
Pathway Enrichment for the 213 Selected Genes. This plot illustrates the biological pathways significantly enriched (FDR*<* 0.05) with the 213 trans-associated genes identified by de-bias SCCA. Here the FDR associated with a pathway represents the probability of that pathway being falsely identified as significantly enriched when it is not. The length of the horizontal bars is proportional to the − log_10_ of the FDR. A longer bar indicates a low FDR, suggesting that the pathway is less likely to be a false positive finding. The horizontal bars are colored by the fold enrichment of the pathways. Fold enrichment is the ratio of the observed overlap between our geneset and a pathway to the expected overlap had there been no enrichment. Higher fold enrichment values signify stronger enrichment of the pathway with the input genes. Thus, the pathways in red are more enriched with our selected genes than the bars in blue.

## 4. Discussion

In this article, we presented a comprehensive pipeline for integrative multi-omics analysis and applied it to identify genes trans-associated with estrogen receptor-related CNAs using data from METABRIC. This study is the first to implement de-biased SCCA, a recent theoretical development in high-dimensional inference, to complex biomedical contexts such as multi-omics analysis, thereby providing a principled analysis pipeline with theoretical guarantees. De-biased SCCA transcends an important limitation of previous SCCA methods by providing asymptotically valid tests for testing whether a given gene is associated with the CNAs and vice versa. Hypothesis tests make false discovery rate control possible via standard FDR control methods (Benjamini and Hochberg, 1995). In the context of trans-association, false discovery rate control is especially crucial because the weaker effect size can potentially lead to spurious detections (Mattioli *et al*., 2020; Alonso *et al*., 2017; Wheeler *et al*., 2019). Moreover, being an SCCA-based approach, our method can aggregate all of the CNA and gene expression information. This is in stark contrast to simpler methods probing only pairwise associations in isolation. In this dataset, our method was able to detect novel trans-associations resulting from weak pairwise associations undetected by linear regression, validating the novel capability of our method. Furthermore, the pipeline based on de-biased SCCA exhibits higher specificity compared to linear regression, which detects most genes and CNAs as associated in this dataset. In contrast, our pipeline leads to a smaller set of genes, many of which have been independently verified to have relevance in breast-cancer-specific associations.

Besides de-biasing, re-sampling offers another potential approach for hypothesis testing with SCCA loadings. To the best of our knowledge, Kessler and Levina (2023) is the only methodological study that systematically explores re-sampling-based tests in this context. However, these tests are based on the ordinary CCA loading vectors (cf. Anderson, 1962) instead of SCCA loadings. Ordinary CCA loadings are not directly computable when the number of variables exceeds the sample size. As such, this method is applicable to scenarios where the sample size is at least as large as the number of variables (*n* ⩾ *p, q*), which is often not the case in CNA-gene expression datasets. For instance, the total number of genes in our module 0 was 5722, far exceeding the sample size of 1904. There is currently no theoretical or empirical evidence that Kessler and Levina (2023)’s method can be extended to high-dimensional settings such as ours. Moreover, re-sampling-based tests are known to encounter subtle pathological issues in high-dimensional penalized estimation frameworks (Chatterjee and Lahiri, 2010, 2011, 2013). Given these considerations, we chose the de-biasing approach over re-sampling for inference using SCCA.

While our analysis is based on the CNA and gene expression data from the METABRIC study, our general pipeline can be extended to any high-dimensional bivariate dataset. However, several considerations should be taken into account during such an extension. First, to screen for CNAs relevant to cancer, we used the ER status as the response variable in our supervised variable selection procedure. This screening step may pose challenges if suitable response variables are unavailable. However, this step is important because it reduces noise and computational burden. At the current state of de-biased SCCA, scaling to unscreened datasets where *p* ∼ 10, 000 is unfeasible. In such cases, unsupervised clustering algorithms may be utilized. Second, the number of selected CNAs, i.e., *p*, is 500 in our analysis, where gene module sizes vary between a few hundred to *q* = 5000. If *p* or *q* are larger, our simulations indicate that the de-biased SCCA algorithm may become conservative. This behavior is expected in high-dimensional tests as the signal-to-noise ratio diminishes with increasing dimensionality (Janková and van de Geer, 2021; Vershynin, 2018). Consequently, the de-biased SCCA method becomes conservative to avoid false positives. Moreover, the initial SCCA-MZ parameter becomes exceedingly sparse in such scenarios. In such cases, rather than using Mai and Zhang (2019)’s SCCA-MZ tuning algorithm (Algorithm **?** in Supplement E) to select the penalty parameter for the initial SCCA-MZ estimator, it may be preferable to choose a smaller penalty parameter. A smaller penalty parameter yields a less sparse preliminary estimator to start with. In our gene module 0, this adjustment ultimately resulted in a slightly larger number of detections (Section 3.3).

Expanding our framework from two to multiple variables via multi-set CCA techniques is a possible avenue for future exploration (Rodosthenous *et al*., 2020; Hu *et al*., 2017; Dashtestani *et al*., 2022). While sparse multi-set CCA has primarily been employed in neuroscience applications, it holds significant promise for genetic data analysis in the near future (Hu *et al*., 2016; Huopaniemi *et al*., 2010; Hu *et al*., 2017; Dutta *et al*., 2014). Finally, the estimation of the preliminary SCCA estimator and the inversion of the Hessian are the current computational bottlenecks of the de-biased SCCA algorithm. These are currently accomplished by the SCCA-MZ estimator and the nodewise lasso algorithm, respectively. These estimators may be replaceable when computationally faster viable alternatives become available, improving the scalability of our method.

In conclusion, our pipeline offers a blueprint for researchers seeking to identify associations among high-dimensional variable sets while simultaneously maintaining principled control over false discovery rates. The ability to aggregate information across all variables makes our method particularly effective at uncovering subtle associations within real-world datasets, especially in biomedical settings. As demonstrated, this approach can reveal important genetic drivers of diseases that might otherwise go undetected due to moderate individual effect sizes, thus generating novel hypotheses for validation in further biological experiments.

## 5. Software

METABRIC data was downloaded from cBioprotal including clinical, genomic and gene-expression data, which had been pre-processed as described previously (Cerami *et al*., 2012). MRMR was implemented using the R package `mRMRe`. For implementing SCCA-MZ, we used the R code provided by Mai and Zhang (2019) with the initiation method “sparse”. To avoid multicollinearity among the CNAs, we added a small amount of Gaussian noise (mean zero, sd = 10^−4^) to the CNA data following O’Driscoll and Ramirez (2016). For gene clustering in METABRIC data, we used the R package WGCNA by Langfelder and Horvath (2008) to perform the `WGCNA` analysis. For de-biasing the SCCA-MZ estimators, we used the R package `de.bias.cca` (Huey and Laha, 2021).

R code for replicating the current analysis, together with a sample input data set and complete documentation, is available on Github at https://github.com/nhuey/dbSCCA.

## Supplementary Material

The supplement to this article contains a collection of simulation results, tables, and figures related to our choice of CNA selection algorithm, the choice of the preliminary SCCA estimator, and choosing ranges for our tuning parameters. Additional material in the Supplement includes the SCCA-MZ algorithm and a figure explaining the differential expression of TM6SF1 and MNDA.

## Acknowledgments

Nilanjana Laha’s research has been partially supported by the NSF-DMS grant DMS-2311098. Diptavo Dutta was supported by the Intramural Research Program of the National Cancer Institute, National Institutes of Health, US Department of Health and Human Services. The authors thank Rajarshi Mukherjee for helpful comments.

## Algorithm 1 An algorithm for tuning the de-biasing procedure

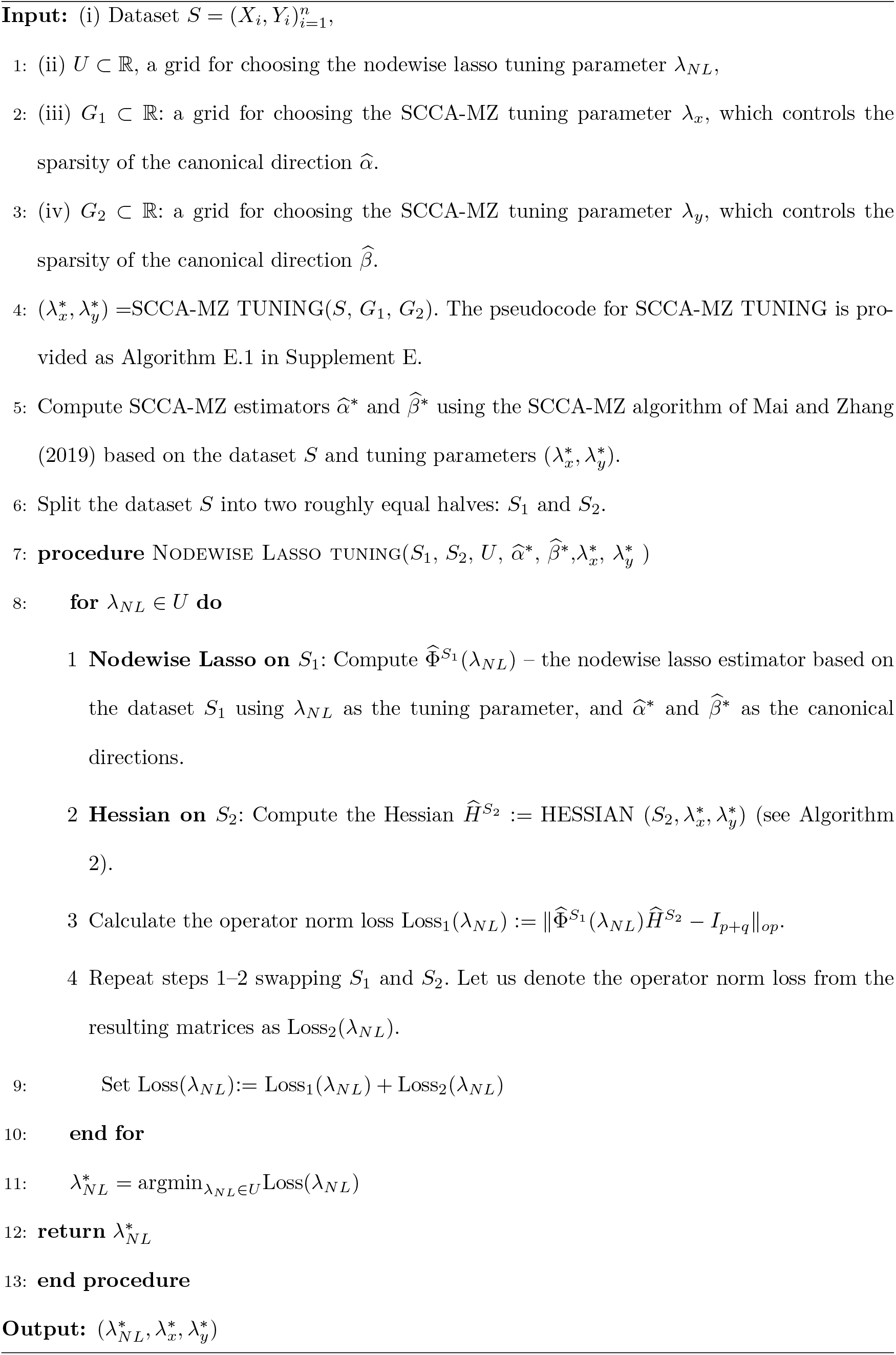

## Algorithm 2 (HESSIAN)

An algorithm for computing the Hessian based on a dataset of size *m*

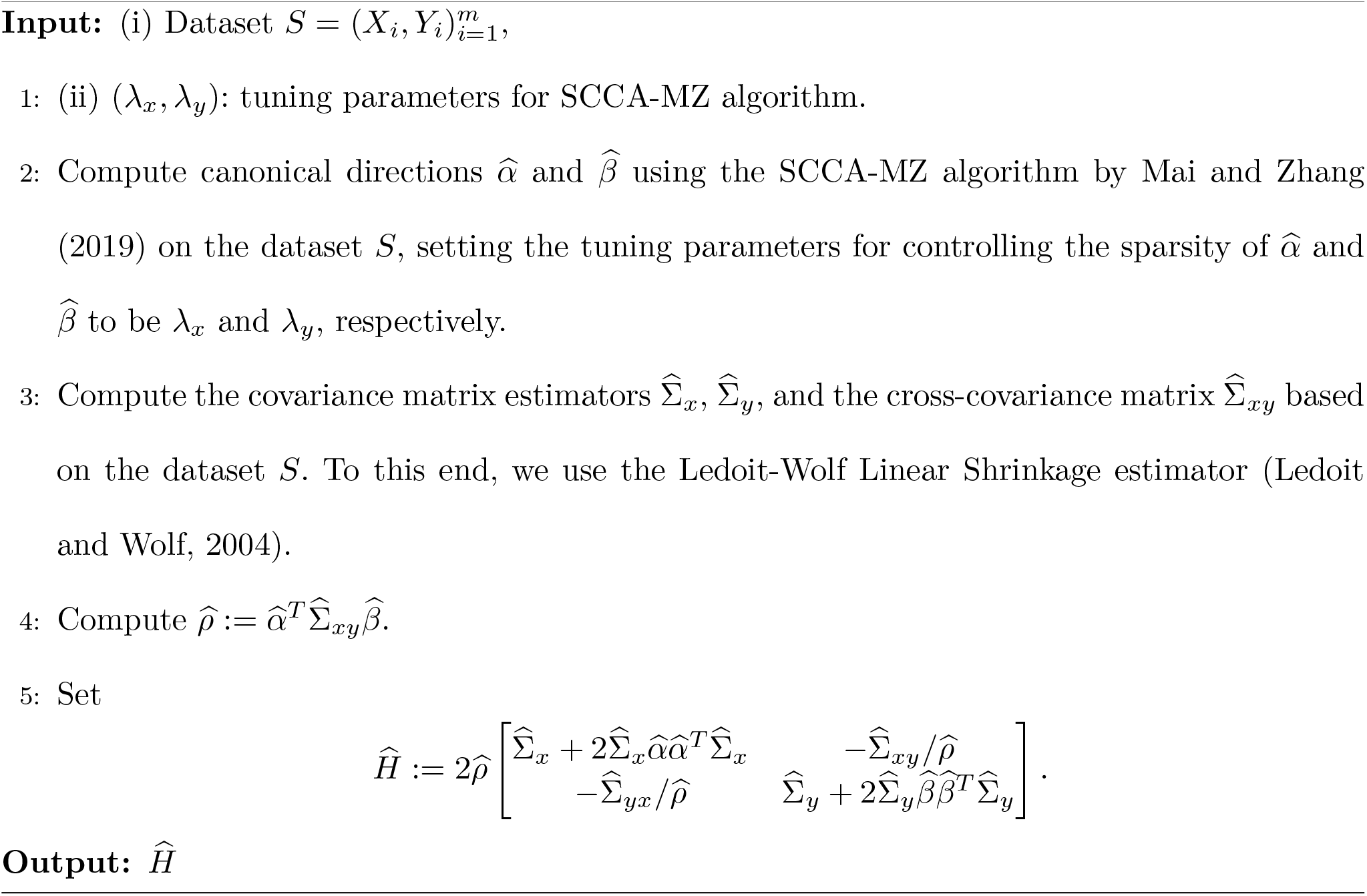

