## Supplementary Material for "De-biased sparse canonical correlation for identifying cancer-related trans-regulated genes"

*Biostatistics* (2024), **0**, 0, *pp.* 1–**??**
doi:10.1093/biostatistics/output

### De-biased sparse canonical correlation for identifying cancer-related trans-regulated genes

NATHAN HUEY[1], DIPTAVO DUTTA[2,*,†], NILANJANA LAHA[3,*,†],
[1] *Mass General Brigham, Mass Eye and Ear Hospital, Boston, MA, USA*
[2] *Integrative Tumor Epidemiology Branch,Division of Cancer Epidemiology and Genetics, National Cancer Institute, Rockville, USA*

[3] *Department of Statistics, Texas A&M University, College Station, TX, USA*

#### A. Justification for the Use of MRMR Algorithm

In this section, we present a comparative study of some feature selection methods on our dataset, which led to the choice of MRMR as our feature selection method. Apart from MRMR, we considered the following three methods for the comparative analysis: Feature Ordering by Conditional Independence (FOCI) (Azadkia and Chatterjee, 2021), RELIEF (Robnik-Šikonja and Kononenko, 2003), and $l_1$-penalized logistic regression (Genç, 2022, cf.).

*To whom correspondence should be addressed.
*† Equal contribution.

© The Author 2024. Published by Oxford University Press. All rights reserved. For permissions, please

*Discussion on the different feature selection methods:* Among the four feature selection methods under consideration, MRMR, FOCI, and RELIEF are filter methods. These types of methods generally take a forward-search-based approach and often produce ranks of the features. We have already discussed the MRMR method in Section 3.1. FOCI uses a measure of conditional dependence designed by Azadkia and Chatterjee (2021), the conditional dependence coefficient (CODEC), to select features using a forward search. The CODEC values can be used to rank the features, with higher values being preferable. Following the stopping rule of Azadkia and Chatterjee (2021), we only use the variables with non-negative CODEC values. We used the R package `FOCI` to implement the FOCI algorithm.

RELIEF is a nearest-neighbor-based method that calculates a score for each feature, ranking the features according to these scores. It is also an iterative method in that the feature scores are calculated using $m$ iterations, where we took $m$ to be 30. In the nearest neighbor step, 10 nearest instances are used. We used the function `attrEval` from R package `CORElearn` to perform RELIEF, taking the evaluation parameter in the function `attrEval` to be "ReliefFequalk".

$l_1$-penalized logistic regression performs a logistic regression using ER status as the response and the CNAs as the predictor variables. An $l_1$ penalty is imposed on the regression coefficients to induce sparsity in the vector of regression coefficients. Only the CNAs with non-zero regression coefficients were selected. This method uses one external tuning parameter, $\lambda$, to control the $l_1$-penalty. To select this tuning parameter, we used a 20% held-out validation set. Using this validation set, the prediction accuracy was calculated for each value of $\lambda$ on a grid of length 100 ranging from 0.0001 to 1. We then chose the $\lambda$ with the highest prediction accuracy for performing the logistic regression.

We use the following procedure for each selection method: we split the dataset using a 3:1 training: test ratio and subsequently generated 40 bootstrapped samples of equal size to the training dataset from the training data. For MRMR and RELIEF, we selected the top 500 CNAs from each bootstrap sample using the scores returned by these methods. The number of selected features obtained by penalized logistic regression on the bootstrap samples was generally less than 500. If they were greater than 500, we chose the top 500 CNAs by ranking them according to the magnitude of the regression coefficients. The selected CNA sets using FOCI were much less than 500 in size in all bootstrap samples. Finally, for each method, we curated a final set of CNAs by combining the CNAs obtained from each bootstrap sample. For MRMR, RELIEF, and logistic regression, we followed the procedure outlined in Section 3.1 to curate a combined CNA set of size 500. However, for FOCI, we simply took the union of all selected CNA sites across the 40 samples. As the individual FOCI-selected CNA sets were small, their union resulted in a set of size 472. It is worth noting that the bootstrapping process described in Section 3.1 for MRMR produced a different CNA set, as it used the entire dataset rather than just the training set.

A.0.1 *Evaluation criteria:* To evaluate the performance of different methods, we considered two metrics: the stability and the prediction of ER status.

*Stability:* Stability is an important criterion in the context of biomarker selection, where it is expected that a desirable algorithm will not drastically modify the top-ranked markers if a few instances were added or deleted (Abeel *and others*, 2010). To assess the stability of a feature selection method, we investigated the pairwise similarity between the 40 CNA sets resulting from the 40 bootstrap samples. To measure this, we used the Kuncheva Index (Kuncheva, 2007), which is a recommended measure in genetic studies for measuring the stability of feature selection methods (Abeel *and others*, 2010). The Kuncheva Index measures the degree of similarity between two sets with the same number of objects in them, i.e., it is a pairwise measurement between two sets of the same cardinality. A higher Kuncheva index corresponds to a higher degree of consistency between the two sets under consideration. Ideally, we would calculate the Kuncheva index between all possible pairs of CNA sets resulting from the 40 bootstrap samples (resulting in 780 indices). This could be done for MRMR and RELIEF because they always selected equal-sized CNA sets, i.e. of size 500, from each bootstrap sample. However, if two sets are of unequal size, the Kuncheva index between them is undefined. Therefore, for methods such as FOCI and logistic regression all possible pairs of CNA sets could not be considered because the selected CNA sets can be different sizes. The Kuncheva index was thus only calculated for those CNA sets with the same size for these methods. A feature selection method is considered stable if the pairwise Kuncheva indices are high. To this end, we considered the average Kuncheva index as well as the overall distribution of the Kuncheva indices for each method.

*Association with ER status:* To assess the association of selected CNAs with the ER status, we evaluated the predictive performance of the final CNA selection sets from different methods. Since ER status can take only two values: $+1$ or $-1$, this prediction can be done by any binary classification algorithm using the selected CNAs as the features. A set of CNAs is considered to be superior if the classifiers exhibit better predictive performance when using these CNAs as predictors. We chose random forests and support vector machines (SVM) as the classification methods. Both these methods are widely used in classification, and have been used for evaluating supervised feature selection methods in existing genetic studies (Pham *and others*, 2017; Mucaki *and others*, 2016). Performance comparison was based on AUC (area under the ROC curve), a standard metric in genetic studies for assessing prediction performance (Abeel *and others*, 2010; Cheng *and others*, 2021; Mucaki *and others*, 2016). A higher AUC value indicates better predictive performance. For random forest, we used 500 trees with 22 variables at each split for all methods except for FOCI, which used 21 variables. For SVM, a cost of 5 and the kernel (sigmoid, radial, or linear) providing the highest AUC were used. For all methods, the radial kernel led to the highest AUC in our data.

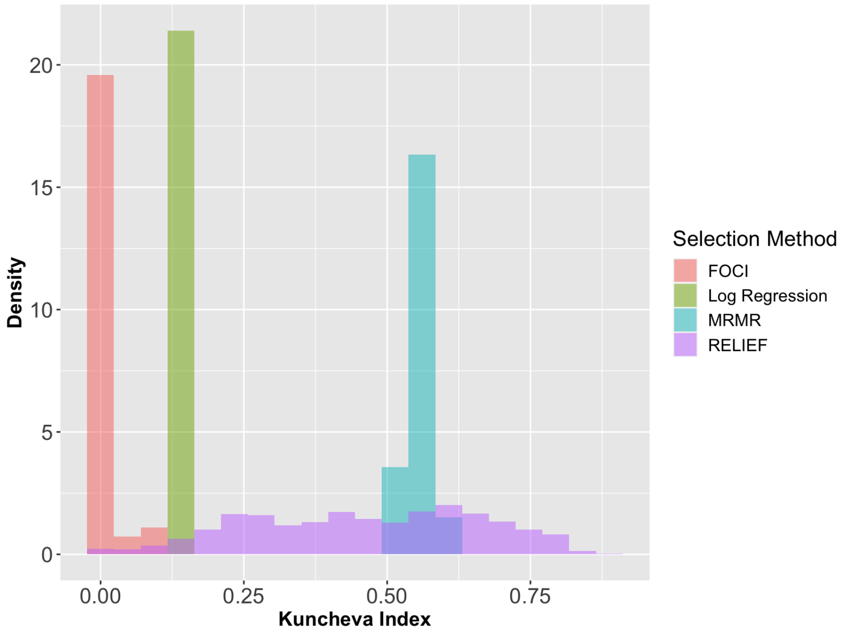


Fig. A.1. **Kuncheva Index Plot for the Feature Selection Algorithms in Section A.** The Kuncheva indices are calculated using all same-sized pairs of CNA selection sets based on the 40 bootstrap samples. Since FOCI and penalized logistic regression generally yield selection sets of unequal sizes, only 2 and 59 indices could be calculated for them, respectively. Both RELIEF and MRMR produced 780 Kuncheva indices, covering all possible pairs. The Kuncheva indices for RELIEF exhibit higher variance, with many values falling below 0.50, the threshold suggested by Kuncheva (2007). All Kuncheva indices for penalized logistic regression and FOCI are below 0.50. FOCI's Kuncheva indices are generally very small. MRMR is the only method with most indices greater than 0.50

A.0.2 *Result:* Table A.1 presents the AUC values for the four methods. MRMR achieves the highest AUC for the random forest classifier, while FOCI and penalized logistic regression demonstrate the highest AUC for the SVM classifier (Table A.1). However, the differences in AUC among these three methods are marginal, not exceeding 0.03 unit for either classifier. Hence, we conclude that FOCI, MRMR, and penalized logistic regression offer comparable prediction accuracy on this dataset. In contrast, RELIEF consistently yields substantially lower AUC compared to the other methods. Figure A.1 shows the distribution of the Kuncheva indices for the four methods based on the bootstrapped data sets. Recall that the Kuncheva index could not be computed for all 780 pairs of selection sets for FOCI and logistic regression. Therefore, the distribution in Figure A.1 considers only the Kuncheva indices that could be calculated for these two methods. Figure A.1 indicates that the Kuncheva indices were generally higher for MRMR compared to the other methods. Kuncheva (2007) considers a feature selection procedure with average Kuncheva index greater than 0.5 to be stable. The average Kutcheva index was 0.556 for MRMR, and 0.463 for RELIEF. The average of the 59 (8% of all possible had all sets been the same size) indices for FOCI was 0.006. The two indices that could be calculated for penalized logistic regression were 0.145 and 0.117. Thus, the Kuncheva indices of MRMR suggest it to be a fairly stable method in terms of CNA site selection. Such guarantees were unavailable for the other methods using Kuncheva indices.

Therefore, our analysis indicates that MRMR offers satisfactory predictive performance with acceptable stability in terms of CNA set selection in our dataset. FOCI and penalized logistic regression yield CNA sets with similar predictive performance, but may lack the desired stability. In view of the above, we chose MRMR as the CNA selection method in our analysis.

Table A.1. **AUC values of different feature selection methods.** Presents the AUC values resulting from classification using the ER status as the class label. The classifications were performed on the test data, using Random Forest and SVM (with radial basis) as the classifiers. For each method, the features in this classification task were this method's final CNA selection sets obtained using the training data. A higher AUC value indicates that the method selected CNAs related to ER status.

| **Methods** | **Random Forest** | **SVM** |
|---|---|---|
| FOCI | 0.763 | 0.792 |
| MRMR | 0.773 | 0.763 |
| RELIEF | 0.578 | 0.545 |
| Penalized logistic regression | 0.768 | 0.793 |

#### B. Choice of Initial SCCA Estimates

##### B.1 *More details on SCCA-MZ*

From Mai and Zhang (2019), it follows that if $\alpha_0$ and $\beta_0$ satisfy some standard sparsity conditions, SCCA-MZ estimators consistently estimate them. In other words, if most CNAs and gene expressions are unassociated, then the SCCA-MZ estimators will be consistent. According to Laha *and others* (2023), the consistency of the preliminary SCCA estimator is a desirable property for the success of the subsequent de-biasing step (Laha *and others*, 2023). Laha *and others* (2023)'s theoretical proof for the asymptotic normality of the de-biased estimators requires the preliminary SCCA estimators to satisfy two more technical conditions. Specifically, it requires the preliminary SCCA estimators to be $\ell_1$ and $\ell_2$ consistent, which means the respective $\ell_1$ and $\ell_2$ distances between $\widehat{\alpha}$, $\widehat{\beta}$, and their population versions should converge to zero in probability (Laha *and others*, 2023). The $\ell_2$ consistency for SCCA-MZ has been proved by Mai and Zhang (2019). However, the $\ell_1$-consistency results for SCCA-MZ have not been explicitly demonstrated.

The SCCA-MZ estimator is an $\ell_1$-constrained estimator because its sparsity is induced by an $\ell_1$ penalty. $\ell_1$-constrained estimators that are $\ell_2$-consistent, are generally also $\ell_1$-consistent, e.g., Gao *and others* (2017)'s COLAR and Janková and van de Geer (2021)'s sparse principal component estimator. Perhaps the best-known example of such an estimator is the standard Lasso under standard regularity conditions (Bühlmann *and others*, 2014). Secondly, the simulation studies in Laha and Mukherjee (2022) indicate that Mai and Zhang (2019)'s method has impressive performance in terms of detecting associations in synthetic Gaussian datasets. Finally, in the course of our own simulations, we have seen that Mai and Zhang (2019)'s method, followed by de-sparsifying leads to qqplots and histograms that suggest Gaussianity of the estimates. This indicates that the theory for de-biasing is likely met by SCCA-MZ estimators, although the demonstration of this fact is beyond the current scope of this work.

##### B.2 *Comparison between SCCA-MZ and COLAR:*

Both COLAR and SCCA-MZ are SCCA estimators proven successful in controlled settings (Mai and Zhang, 2019; Gao *and others*, 2017). This section presents a small-scale study comparing the performance of COLAR and SCCA-MZ as preliminary estimators for de-biasing procedures in noisy biomedical datasets. We used a subset of the METABRIC data to conduct this study. We kept the set of 500 CNAs selected by MRMR (Section 3.1), but considered smaller subsets of genes that are already known to be implicated in cancer. To this end, we used two gene-subsets. The first set is a list of 154 genes, given by Bertucci *and others* (2006), that are implicated in a rare type of breast cancer called medullary breast cancer (Bertucci *and others*, 2006). We will refer to the resulting subset of our data as the "medullary breast cancer dataset". The second geneset comprises 167 genes belonging to a geneset found by Curtis *and others* (2012) to be frequently mutated in breast cancer. We call the resulting subset of our dataset the "PCAWG dataset" naming it after Curtis *and others* (2012)'s study which was based on the Pan-Cancer Analysis of Whole Genomes (PCAWG) Consortium of the International Cancer Genome Consortium and The Cancer Genome Atlas.

We performed the de-biasing procedure using both SCCA-MZ and COLAR on these two datasets as well as a shuffled version of them. The shuffling was performed by randomly permuting the rows of the CNA data matrix to break any associations between CNA and gene expression. The data shuffling is supposed to weaken the association between the genes and the CNAs. If the patients are unrelated, then one would expect the genes and CNAs to be uncorrelated in the shuffled dataset, representing the null case: $\alpha_i = 0$ and $\beta_j = 0$ for all $i \in \{1, \ldots, p\}$ and all $j \in \{1, \ldots, q\}$. Therefore, rejections of the tests $H_0 : \alpha_i = 0$ and $H_0 : \beta_i = 0$ in the shuffled data reflect false positives. Hence, the method with fewer rejections in the shuffled dataset will be preferred.

We consider the $i$th CNA to be "detected" if the hypothesis test $H_0 : \alpha_i = 0$ corresponding to this CNA is rejected, and similarly for the $j$th gene. To correct for potential false positives, we used a conservative p-value cutoff $2.5 \times 10^{-6}$ in the above tests. COLAR was implemented using the MATLAB code provided by Gao *and others* (2017) with the default parameters. The tuning parameters for SCCA-MZ were chosen using cross-validation. Table B.1 displays the number of detected CNAs and genes in both datasets and their shuffled versions.

Applying de-biasing with COLAR increased the number of detected CNAs from 142 to 154 in the medullary breast cancer dataset after shuffling. In the PCAWG dataset, although the number of detected CNAs did not increase, de-biased COLAR still detected 83 CNAs after shuffling, reflecting only a 32% reduction due to shuffling. This observation hints that the COLAR-based de-biasing method can mistakenly assume some CNAs to be associated with the genes when there is no association at all, indicating an inclination towards false positives while analyzing CNA-gene data. In contrast, the SCCA-MZ-based de-biasing algorithm detected only one CNA in the shuffled medullary breast cancer dataset and four CNAs in the shuffled PCAWG dataset. These numbers are much smaller compared to SCCA-MZ's detections in the unshuffled datasets, which were 187 and 208, respectively. Neither method detected many genes in either the original or shuffled datasets. However, de-biased COLAR still detected one gene in the shuffled PCAWG dataset, whereas de-biased SCCA-MZ did not detect any gene in either of the shuffled datasets.

In view of SCCA-MZ's superiority in preserving type I error in the subsets of the METABRIC data, we selected it as the preliminary SCCA estimator for our analysis.

Table B.1. **Size of selected CNA and gene sets for the Medullary Breast Cancer and PCAWG datasets before and after shuffling.** Shuffling was performed by randomly permuting the rows of the CNA data matrix to destroy any association between CNAs and genes. Both SCCA-MZ and COLAR were followed by the de-biasing method to enable hypothesis testing. A p-value cut off of $2.5 \times 10^{-6}$, common in genetics literature, was used in the hypothesis tests as protection from potential false positives. However, the number of detections by de-biased COLAR did not decrease as much compared to SCCA-MZ after shuffling.

| **Datasets** | **Medullary Breast Cancer** | | **PCAWG** | |
|---|---|---|---|---|
| | **CNA (p = 500)** | **Genes (q = 154)** | **CNA (p = 500)** | **Genes (q = 167)** |
| **SCCA-MZ** | 187 | 2 | 208 | 6 |
| **COLAR** | 142 | 18 | 122 | 8 |
| **SCCA-MZ (shuffled)** | 1 | 0 | 4 | 0 |
| **COLAR (shuffled)** | 154 | 0 | 83 | 1 |

#### C. Choosing the grid for SCCA-MZ tuning parameters

To choose the grid for tuning the SCCA-MZ parameters $\lambda_x$ and $\lambda_y$, we conducted a small-scale simulation study with the following data-generation scheme, motivated by parameters from the METABRIC data. We considered the parameter combination $(n, p, q) = (1900, 500, 150)$. The data matrices $X$ and $Y$ were generated according to the sparse inverse setting outlined in Section 5 of Laha *and others* (2023), where the population-level covariance matrices $\Sigma_x$ and $\Sigma_y$ are sparse inverse matrices (see Laha *and others*, 2023, for more details) and $\Sigma_{xy} = \rho_0 \Sigma_x \alpha_0 \beta_0^T \Sigma_y$. We selected the vectors $\alpha_0$ and $\beta_0$ according to the sparse inverse setting of Section 5 of Laha *and others* (2023), with their sparsity (number of non-zero elements) set to 150 and 10, respectively. We set $\rho_0 = 0.7$, a conservative choice given that the SCCA-MZ estimates of the canonical correlations across the 6 modules fell within the range of $[0.83, 0.92]$. This decision was motivated by our observation of upward bias in the SCCA-MZ estimators of canonical correlations during simulations. We implemented SCCA-MZ with the default parameters in the code provided by Mai and Zhang (2019). For initializing SCCA-MZ, we chose a random initialization where each element of $\alpha_0$ and $\beta_0$ were sampled from a standard Gaussian random vector.

We generated 100 samples using the above simulation scheme. SCCA-MZ often did not run under the current simulation setting when $\lambda_x$ and $\lambda_y$ were greater than 0.05. Therefore, we considered a $50 \times 50$ grid of equally spaced tuning parameter pairs $(\lambda_x, \lambda_y)$ ranging from 0 to 0.05. We observed that the combined $l_2$-error of the estimated canonical directions $\widehat{\alpha}$ and $\widehat{\beta}$ ranged between 2.903495 and 3.3 on this grid. In the final data application, we chose the grid to be [0, 0.10] to be conservative.

#### D. Choosing the grid for nodewise lasso tuning

In this section, we provide the rationale behind the selection of the grid for tuning the nodewise lasso parameter $\lambda_{NL}$, as discussed in Section 3.3. We conducted a pilot study generating $N = 100$ datasets using the data generation scheme outlined in Section C, with two modifications. Firstly, we adjusted the $(n, p, q)$ combination to $(100, 50, 50)$ in this pilot study. These values for $p$ and $q$ are smaller compared to those in Section C. This adjustment accounts for the higher computational intensity of nodewise lasso, particularly for larger values of $p$ and $q$. Secondly, we set the number of non-zero elements in $\alpha_0$ and $\beta_0$ to be 3. We considered a grid of 50 equally spaced points ranging from 0.01 to 10 for selecting $\lambda_{NL}$. For $\lambda_{NL} \geqslant 10$, the nodewise lasso algorithm did not run to completion in many replications.

Now we outline our simulation procedure. For each tuning parameter $\lambda_{NL}$, and each replication $i = 1, \ldots, 100$, we performed the following:

1. We generated data according to the above-mentioned simulation scheme. Then we computed the SCCA-MZ estimators using Mai and Zhang (2019)'s algorithm, using default parameters and the random initiation described in Section 2.5. At this step, we selected the SCCA tuning parameters $(\lambda_x, \lambda_y)$ using Algorithm E.1. The tuning parameters were chosen from a $10 \times 10$ grid of 10 equally spaced tuning parameters ranging from 0 to 0.25.

2. We split the data according to 3:1 training: test ratio.

3. We computed the nodewise lasso estimator $\widehat{\Phi}_i(\lambda_{NL})$ using the training data and the SCCA-MZ estimators derived at the first step.

4. We computed the hessian estimator $\widehat{H}_i$ based on the validation set using Algorithm 2. Algorithm 2 requires two SCCA-MZ tuning parameters as input, which were chosen using the procedure mentioned in step 1 on the validation set.

5. The $i$th replication's operator norm loss with respect to the estimated hessian $\widehat{H}_i$ was computed as follows:

$$\text{Loss}_i(\lambda_{NL}) = \|\widehat{\Phi}_i(\lambda_{NL})\widehat{H}_i - I_{p+q}\|_{op}. \tag{D.1}$$

A version of the above loss has been used to find the optimal tuning parameter during the nodewise lasso tuning procedure in our main analysis (see Section 2.5). In this simulation study, we also computed the operator norm loss with respect to the true hessian $H_0$ (see Laha *and others*, 2023 for its expression), given by

$$\text{Loss}_i^0(\lambda_{NL}) = \|\widehat{\Phi}_i(\lambda_{NL})H_0 - I_{p+q}\|_{op},$$

to assess the actual deviation of $\widehat{\Phi}_i(\lambda_{NL})$ from $H_0^{-1}$.

We averaged the losses across all completed replications for each $\lambda_{NL}$, resulting in two average losses: the estimated operator norm loss (uses the estimated hessian) and the true operator norm loss (uses the true hessian). We will denote these losses by $\text{Loss}(\lambda_{NL})$ and $\text{Loss}^0(\lambda_{NL})$, respectively. Figure D.1 plots these two losses as a function of $\lambda_{NL}$. The estimated operator norm loss was much higher than the true operator norm loss, which is unsurprising due to the large operator norm errors in high-dimensional matrix estimation (Vershynin, 2010). To visualize both losses on the same plot, we scaled the estimated operator norm loss by $(p+q)^{-1}$ in Figure D.1.

Figure D.1 highlights that the estimated operator norm loss continues to decrease with $\lambda_{NL}$. The true operator norm loss is also non-increasing with $\lambda_{NL}$ but changes negligibly for larger values. Therefore, it seems reasonable to use the estimated operator norm loss to guide tuning parameter selection from the chosen range. Due to the low variability of the true operator norm loss for $\lambda_{NL} \geqslant 5$ and the computational expense of the nodewise lasso algorithm, we opted for a coarser grid of length 5 over the range $[0, 10]$ for tuning $\lambda_{NL}$ in our primary analysis.

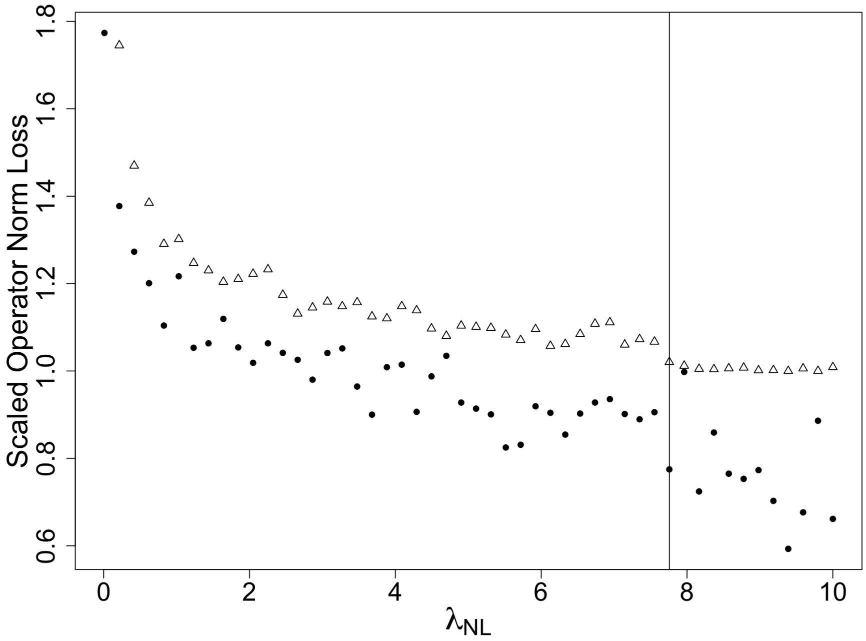


Fig. D.1. **Operator norm loss vs nodewise lasso tuning parameter $\lambda_{NL}$.** The solid dots display the scaled (by $1/(p+q)$) estimated operator norm loss, while the triangles display the true operator norm loss of the inverted hessian estimator $\widehat{\Phi}(\lambda_{NL})$. The true operator norm loss captures the departure of $\widehat{\Phi}(\lambda_{NL})$ from the true inverted hessian. The estimated operator norm loss has been used to choose the optimal $\lambda_{NL}$ in Algorithm 1, measures this departure with respect to an estimated hessian. We used 100 replications to calculate these losses. The vertical line indicates $\lambda_{NL} = 7.76$, beyond which fewer than 50% of the replications completed.

#### E. The SCCA-MZ algorithm

**Algorithm E.1 SCCA-MZ TUNING:** An algorithm by Mai and Zhang (2019) for tuning the SCCA-MZ estimators

**Require:** (i) Dataset $S = (X_i, Y_i)_{i=1}^m$,

(ii) $G_1 \subset \mathbb{R}$: a grid for choosing the SCCA-MZ tuning parameter $\lambda_x$.

(iii) $G_2 \subset \mathbb{R}$: a grid for choosing the SCCA-MZ tuning parameter $\lambda_y$.

**for** $\lambda_x \in G_1$ **do**

  **for** $\lambda_y \in G_2$ **do**

    **Five-fold cross validation:**

Split the dataset $S$ into five equal folds: $F_1$, …, $F_5$.

    **for** $i \in \{1, \ldots, 5\}$ **do**

1. Compute the sample cross-covariance matrix $\widehat{\Sigma}_{xy}^{(i)}$ based on the fold $F_i$.
2. Compute the SCCA-MZ estimators $\widehat{\alpha}^{(i)}$ and $\widehat{\beta}^{(i)}$ based on the remaining folds. To this end, use the tuning parameters $\lambda_x$ and $\lambda_y$ to control the sparsity of $\widehat{\alpha}^{(i)}$ and $\widehat{\beta}^{(i)}$, respectively.
3. Calculate $\widehat{\rho}^{(i)} = (\widehat{\alpha}^{(i)})^T \widehat{\Sigma}_{xy}^{(i)} \widehat{\beta}^{(i)}$.

    **end for**

    Set $\widehat{\rho}(\lambda_x, \lambda_y)$ to be the average of $\{\widehat{\rho}^{(1)}, \ldots, \widehat{\rho}^{(5)}\}$.

  **end for**

**end for**

$(\lambda_x^*, \lambda_y^*) = \text{argmax}_{(\lambda_x, \lambda_y) \in G_1 \times G_2} \widehat{\rho}(\lambda_x, \lambda_y)$

**Ensure:** $(\lambda_x^*, \lambda_y^*)$

#### F. Supplementary Figure

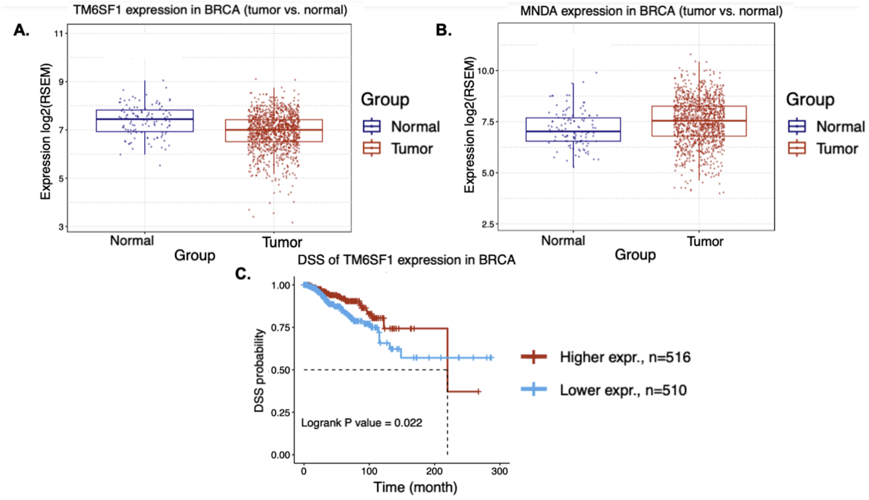


Fig. F.1. **Differential expression of *TM6SF1* and *MNDA*.** **A**. Boxplots displaying the gene expression (measured in RSEM units) of *TM6SF1* in normal and tumor tissues of TCGA-Breast Invasive Carcinoma (BRCA) data. **B**. Boxplots displaying the gene expression (measured in RSEM units) of *MNDA* in normal and tumor tissues of TCGA-BRCA data. These plots suggest higher expression of *MNDA* in tumor cells. **C**. Kaplan-Meier plot for disease-specific survival (DSS) comparing high and low expressions of *TM6SF1* in the TCGA-BRCA dataset. Gene expression of *TM6SF1* was dichotomized at the median across all samples.

[*Received August 1, 2010; revised October 1, 2010; accepted for publication November 1, 2010*]
